# Comparison between cultivation and sequencing based approaches for microbiota analysis in swabs and biopsies of chronic wounds

**DOI:** 10.1101/2020.09.08.288779

**Authors:** Aleksander Mahnic, Vesna Breznik, Maja Bombek Ihan, Maja Rupnik

## Abstract

Chronic wounds are a prominent health concern affecting 0.2% of individuals in the Western population. Microbial colonization and the consequent infection contribute significantly to the healing process of chronic wounds. We have compared cultivation and 16S amplicon sequencing (16S-AS) for the characterization of bacterial populations in swabs and biopsy tissues obtained from 45 chronic wounds and analysed metadata for wound-specific and clinical-outcome-associated correlations with bacterial community structure.

Using cultivation approach, we detected a total of 39 bacterial species, on average 2.89 per sample (SD=1.93). Comparison of cultivation results between swabs and biopsy samples showed no significant advantage of one sampling method over the other. 16S-AS was advantageous in comparison to the cultivation approach in case of highly diverse communities, where we could additionally detect numerous obligate and facultative anaerobic bacteria from genera *Anaerococcus, Finegoldia, Porphyromonas, Morganella* and *Providencia*. Based on the community diversity, chronic wound microbiota could be distributed into three groups, however, no correlation between groups and clinical outcome was observed. Clinically estimated presence of biofilm and a larger surface area at the initial visit were most significantly associated with unfavourable clinical outcomes after one-year follow-up visit. *Corynebacterium* was the single most predictive bacterial genus associated with unfavourable clinical outcomes in our study.

## Introduction

Chronic wounds (CWs) are commonly defined as wounds that fail to spontaneously heal in six weeks (1) and are commonly classified into three most prevalent etiological categories: 1) venous valve insufficiency and dependency, 2) lower extremity arterial disease and 3) diabetes (2). They affect approximately 2.21 per 1000 individuals in the Western population, significantly reducing life quality of the patients and representing a costly burden for the health system (3).

Wound healing is a complex process, affected by various systemic and local factors, among which microbial burden is one of the major culprits for non-healing CWs (4). Colonization with bacteria and fungi has been previously linked to different CW specific parameters, temporal dynamics and healing outcomes (5–9). The impaired CW healing is partially associated with biofilm formation, which provides resistance against host defences and antimicrobial therapy (10, 11). The role of individual bacterial species in CW developement and healing process, however, remains largely unclear and results in frequent overuse and ineffectiveness of antimicrobial agents in the treatment of CWs (12, 13).

The diagnosis of CW infection currently relies on a combination of clinical judgement and microbiological cultivation of different specimens: swabs, which are non-invasive and more frequently used, or wound tissue, obtained by a more demanding and invasive biopsy or curettage (14). Recent guidelines for CW biofilms specify tissue biopsies as a gold standard for microbiological diagnostics due to the sampling of both surface and deeper tissue (4).

Molecular approaches enabling the exact and cost effective characterization of mixed microbial populations are slowly being integrated into microbiological diagnostics in general (15) and have been considered for diagnostics of CWs (16). Characterisation of the microbiota in CWs is crucial in order to improve our understanding of its impact on the healing process. It is intuitive to assume that the molecular characterisation, especially high-throughput sequencing, will outperform cultivation-based methods, because of the limitations associated with culturing the slow-growing and anaerobic microorganisms. However, only few studies up to date systematically evaluated the differences between the two approaches (17–19). Previous studies specifically performed on CWs used either sequencing-based methods (20) or cultivation-based methods (14, 21, 22) to compare between the swab and biopsy specimens.

In this study, we used sequencing- and cultivation-based approaches to analyse paired swab and biopsy specimens in order to evaluate different methodologies for characterizing the complex bacterial populations in CWs. Additionally, extensive metadata were used to reveal wound-specific and clinical-outcome-associated correlations with bacterial community structure.

## Methods

### Specimen collection

The study included 45 in- and outpatients with CWs, who were treated in the Department of dermatology and venereal diseases at University Medical Centre Maribor from February to June 2017. The inclusion criteria were: age above 18 years and CWs with duration of more than six weeks. Only one CW per patient was sampled. The majority of CWs were located on the lower legs and were classified into four etiological categories: venous/dependency (n=30), mixed arterial-venous (n=6), diabetic wounds (n=2) and wounds of other etiologies (n=7). The ethical approval was obtained by institutional ethical board committee (UKC-MB-KME-23-13/17).

After obtaining patients’ written informed consents, clinical data relating to the characteristics of the patients (age, gender, body mass index (BMI) and ankle-brachial index) and of the CWs (duration, estimated surface area calculated as length x width of the wound, clinical assessment of biofilm presence and clinical outcome of the wound after one year), were obtained. Although, no generally accepted clinical signs of CW biofilms exist (23), extensive fibrinous slough has been proposed as possible macroscopic clue of CW biofilm (24) and this was also adopted in this study.

All CWs were cleaned with potable warm water, soap (pH 5.5), gauze and a round curette. The swabs were obtained according to the Levine technique, which consists of rotating a swab over one cm^2^ area with sufficient pressure to express fluid from within the CW tissue (22, 25). The swabs were placed in a sterile container with liquid medium and were sent within 24 hours for downstream analysis. Next, 3–4 mm punch biopsies were performed on the same CW area under local anesthesia and the tissue samples were bisected, with half of the sample immediately frozen and stored in liquid nitrogen at −197 °C until NGS, and the other half placed in sterile container and sent within 24 hours for bacterial cultivation.

### Cultivation-based analysis of samples

Swabs were re-suspended in the physiological solution. Suspension was divided in two aliquots, one of them was used for DNA isolation and sequencing, the other was initially inoculated into thioglycolate enrichment. This was subsequently cultured on the blood agar and selective media for Gram negative bacteria. Plates were incubated either at 5% CO_2_ or at aerobic atmosphere.

Tissue biopsies were initially homogenized in physiological solution (Millimix 20, DOMEL). The homogenate (100 µL) was transferred into thioglycolate broth and into cooked meat broth. After 24h, enrichment cultures were inoculated on the same media and under same conditions as described above for swabs. Additionally, anaerobic cultivation was performed on COH and Schaedler agar. Colonies were isolated in pure culture and identified with Maldi Biotyper (Bruker Daltonik).

### 16S metagenomic sequencing

The tissue biopsies and swab sample residues (in physiological solution) were stored at −80 °C until molecular diagnostics. Total DNA was extracted with QIAamp DNA Mini kit (QIAGEN) with a modified protocol. Pellets were re-suspended in 360 μL of ATL buffer and homogenized (SeptiFast tubes (Roche), MagnaLyser (Roche), 7000 rpm, 70 s). Afterwards, 40 μL of proteinase K was added and the suspension was incubated at 55 °C for 1h. Next, 200 μL of AL buffer was added, followed by an incubation at 70 °C for 30 min. After the addition of 200 μL of 96-100% ethanol, we transferred the content into column tubes and the subsequent steps followed the protocol provided in QIAamp DNA Mini kit. Extracted DNA was stored at −80 °C until further use.

We sequenced the V3V4 variable region of the 16S rRNA gene. Libraries were prepared according to the 16S Metagenomic Sequencing Library Preparation (Illumina) protocol using the primer pair Bakt_341F (5’-CCTACGGGNGGCWGCAG-3’) – Bakt_805R (5’-GACTACHVGGGTATCTAATCC-3’), approximately 460 bp fragment length (26). Library quality was checked with Bioanalyzer High Sensitivity DNA Assay. Sequencing was performed on the Illumina MiSeq platform (2×300 bp, 5 % PhiX).

### Data availability

Sequence reads are available via SRA with BioProject number PRJNA659504.

### Sequence data analysis and statistics

Quality filtering was performed using mothur software (27) with parameters as recommended by Kozich et al. (28). Alignment was performed using Silva reference base (Release 123). Chimeras were identified using mothur implemented UCHIME algorithm. Taxonomy was inferred with the RDP training set (v.12) (0.80 bootstrap value). We obtained a total of 7576427 reads (min: 14453, max: 81825, average per sample: 39460.56).

A negative control (dH_2_O) was included in every DNA isolation batch (14 samples), totalling 5 negative controls. High abundance of contaminants is expected in samples with low bacterial burden such as CWs. To eliminate as many contaminants as possible from our dataset we implemented the following procedure: for each OTU we selected the negative control with the highest number of reads (N_max_). Respective OTU was then removed from all samples with less than 5 x N_max_ reads.

Statistical analysis was done in mothur (alpha and beta diversity, Bray-Curtis dissimilarity) and in R using packages ‘vegan’ and ‘ggplot2’. Analysis of sequencing reads at species taxonomical level for *Corynebacterium* genus was performed with Oligotyping tool (29) and BLAST (https://blast.ncbi.nlm.nih.gov/Blast.cgi; 5.5.2020).

## Results

### Cultivation-based analysis of swabs and biopsy specimens

Swabs and biopsy specimens were collected from 45 CWs. Using cultivation approach, we detected a total of 39 different bacterial species, on average 2.89 species per sample (SD=1.93). The average number of detected species per sample did not differ between biopsy and swab specimens (pairwise t-test, p=0.93). The structure of bacterial population however showed only partial congruence (Fig. 1, Table S1). In 58.4% of cases, a respective bacterial species was detected in both specimens of the same CW, denoted hereinafter as a match. Highly prevalent bacterial species were more likely to match in swab/biopsy pairs of samples. For instance, when bacterial species was detected in at least five CW samples, we observed a 68% matching rate compared to the 37% matching rate in case of species that were detected in less than five CW specimens (Fisher exact test, p=0.027). In 41.6% of paired swab/biopsy samples we observed a miss-match, i.e. cases where a bacterial species was detected in only one of the paired swab/biopsy samples. Similar rates of miss-matches were observed for swabs (20.5%) and biopsy samples (21.1%) (Fisher exact test, p=1.000).

**Figure 1:**
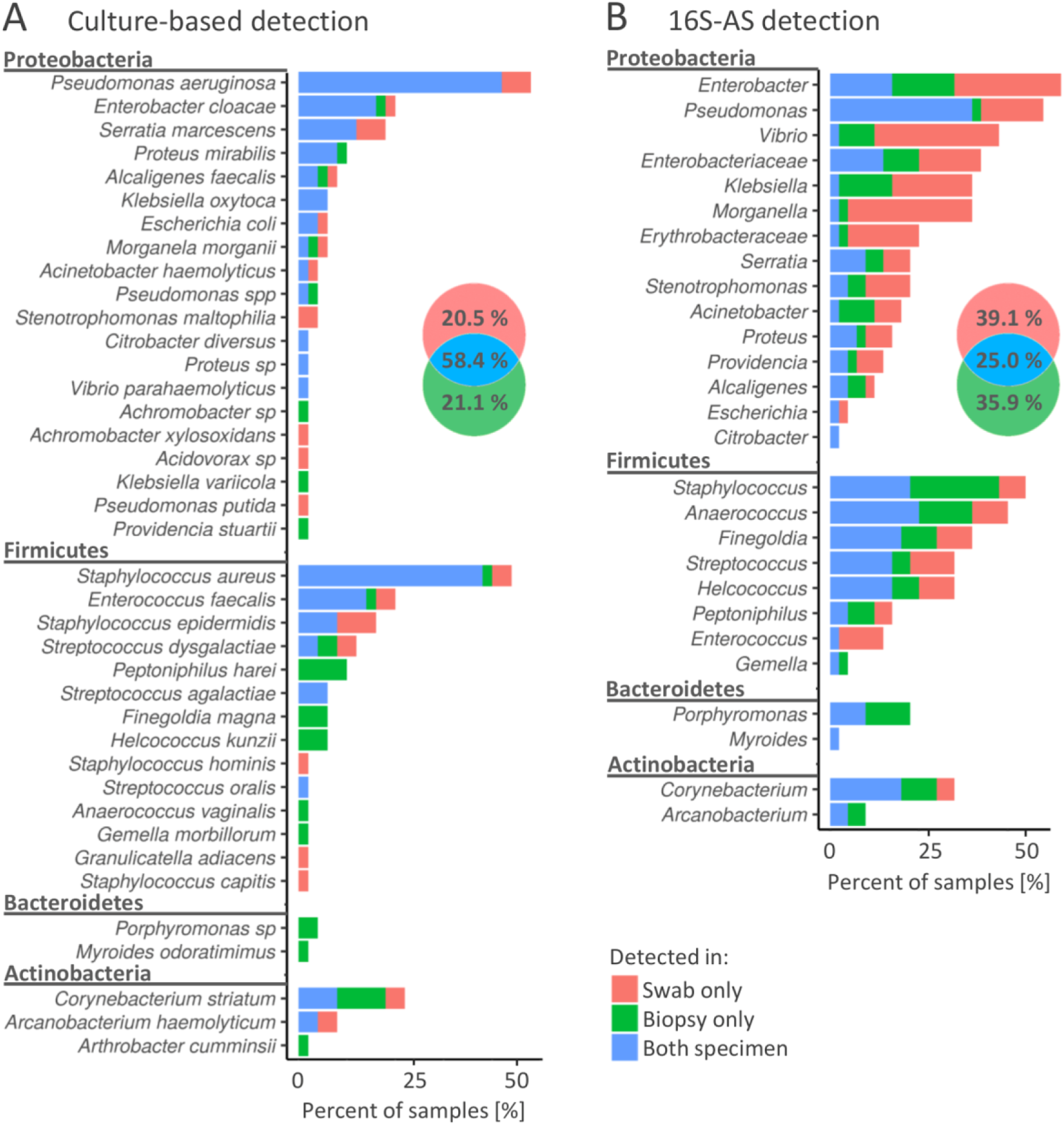
Concordance in the detection of bacterial species (cultivation) or genera (16S-AS) in swabs and biopsy specimens obtained from chronic wounds. Colours in the frequency histograms denote the percentages of samples according to the matching/non-matching detections between swab/biopsy sample pairs of the same CW. Results are presented for cultivation-based approach (A) and 16S-AS (B), separately. Venn diagrams show cumulative percentages for all detected bacterial groups. Note that panel A (cultivation) represent diversity at the species taxonomic level while panel B (16S-AS) at the genus taxonomic level, therefore the direct comparison of detectable diversity between methods is not possible in this figure.

Cultivable bacterial community was represented by four bacterial phyla, of which *Proteobacteria* and *Firmicutes* showed the highest diversity and prevalence (Fig. 1A). *Pseudomonas aeruginosa* and *Staphylococcus aureus* were the most prevalent species and were detected in 53% (24/45) and 49% (22/45) of CWs, respectively. In contrast, 43.6% (17/39) of detected species were each detected in only one CW sample (Suppl. Table S1).

When using cultivation approach, biased detection between specimens was most notable among the less prevalent bacterial species. For example, *Peptoniphilus harei* (n=5), *Finegoldia magna* (n=3), *Helcoccus kunzii* (n=3) and *Porphyromononas sp*. (n=2) were detected several times, but only in biopsy samples. On the other hand, *Stenotrophomonas maltophilia* (n=2) was detected only in swabs (Fig. 1A).

### Sequencing based analysis of swabs and biopsy specimens

Amplicon sequencing of the V3V4 variable region of 16S rRNA gene (16S-AS) yielded 73 operational taxonomic units (OTUs), on average 5.9 per sample (SD=7.1). Similar to cultivation approach, the identified OTUs comprised four bacterial phyla, with *Proteobacteria* and *Firmicutes* dominating the complex communities (Fig. 1B). Most prevalent were representatives from genera *Enterobacter, Pseudomonas* and *Staphylococcus*. 16S-AS analysis revealed culture-approach-associated under-representation of several taxa, mainly *Vibrio, Anaerococcus, Finegoldia* and *Enterobacteriaceae*.

The rate of miss-matches between swabs and biopsy samples was greater when using 16S-AS as compared to the cultivation-based approach (75% vs 41.6%), This is likely a consequence of detecting bacteria, which are present in low bio-burden (Fig. 1B). In both, culture- and 16S-AS-based approach, the detection of *Porphyromonas* was more consistent in biopsy specimens compared to swabs.

### Comparison between 16S amplicon sequencing and cultivation-based approach

By using the sequencing data, CW samples were distributed into three groups based on the number of OTUs that were required to cover 99% of the obtained number of reads. Diversity group A included low diversity samples with a single OTU comprising 99% of all reads; diversity group B required 2 to 4 OTUs to cover 99% of obtained reads while the diversity group C required more than 4 OTUs (Fig. 2, Fig. S1 and S2). Only 51.1% (23/45) swab/biopsy pairs were classified into the same diversity group.

**Figure 2:**
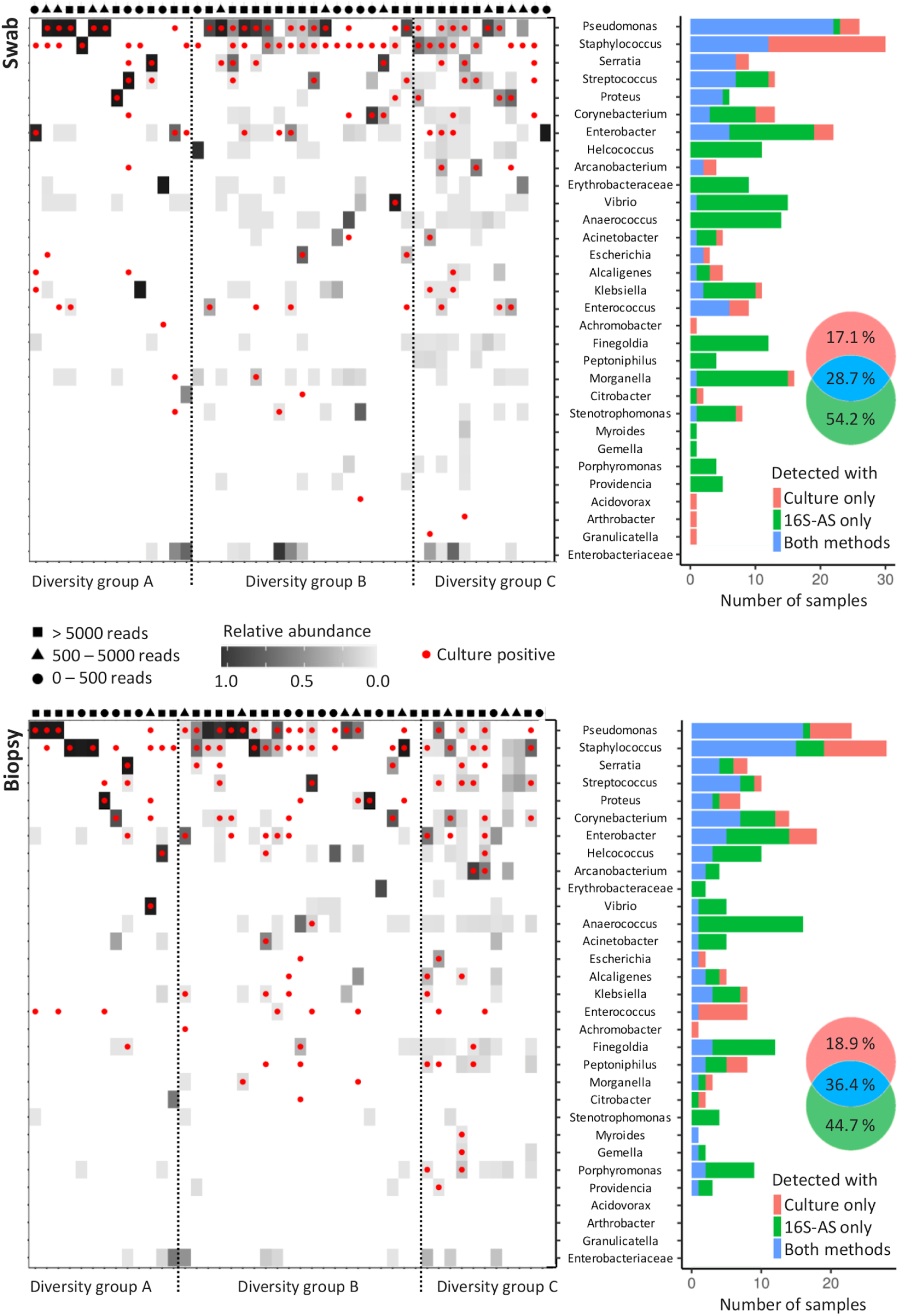
Concordance in the detection of bacterial genera with cultivation and 16S-AS in chronic wounds. Heat-plots show relative abundances of bacterial genera across all analysed CW samples obtained from swabs (top) and biopsies (below). Red dots denote genera, which were also detected with cultivation-based approach. Samples were assigned to three diversity groups (A, B and C), based on the number of operational taxonomic units (OTUs) that were required to cover 99% of total obtained sequencing reads. Diversity groups were not associated with the number of reads obtained per sample, indicated by the symbols shown above the heat-plots approximating the number of reads per sample into three categories. Histograms (right) show the concordance between cultivation-based detection and 16S-AS. Venn diagrams show cumulative percentages for all detected bacterial genera.

The concordance between 16S-AS and cultivation-based detection of bacterial genera depended on the community diversity. Our results indicate that cultivation-based detection underestimates the bacterial richness in highly diverse bacterial communities (diversity group C) while no such bias was observed in case of low diversity communities (diversity group A) (Fig. S3). The concordance between 16S-AS and cultivation-based detection was higher in biopsy samples (36.4% matching rate) compared to the swab samples (28.7% matching rate) (Fisher exact test, p=0.002). Representatives from genera *Pseudomonas, Staphylococcus* and *Enterococcus* were most often detected with cultivation method while absent in 16S-AS data. Moreover, representatives from genera *Achromobacter, Acidovorax, Arthrobacter* and *Granulicatella*, each detected once with cultivation, were not detected with 16S-AS. *Achromobacter* was the only one among these, which was removed from 16S-AS data due to the high number of reads found in the negative controls. On the other hand, several genera were often detected with 16S-AS, but not with cultivation. These included anaerobic representatives from *Anaerococcus, Finegoldia* and *Porphyromonas*; facultative anaerobes *Morganella, Vibrio* and *Providencia*, and aerobes from genera *Erythrobacteriaceae* and *Stenotrophomonas* (Fig. 2).

### Microbiota associations with patient- and wound-specific parameters and clinical outcome

In this study, underlying causes of the CWs included venous insufficiency/dependency, combination of peripheral arterial disease and venous insufficiency, diabetes and other causes (see Materials and methods). Due to the large disproportion in the number of samples of CWs from each etiological category, comparison between them was not possible.

Patient-specific factors (age, gender and body mass index) were not significantly correlated with bacterial community structure neither in swab nor in biopsy specimens (Permutational multivariate analysis of variance (PERMANOVA) using Bray-Curtis distances, p>=0.42; Table S2).

Wound-specific parameters included CW duration, estimated CW surface area, ankle-brachial index, clinical assessment of biofilm and the clinical outcome at the one-year follow-up visit. Complete metadata along with OTU-based community structure is available in Figures S1 and S2 for swabs and biopsy samples, respectively. *Pseudomonas* (OTU1) was the only bacterial group significantly associated with a larger CW surface area, both in biopsy (Spearman’s r=0.55, p<0.001) and swab specimens (Spearman’s r=0.48, p=0.001; Fig. 3A). The CW surface area was also positively correlated with the bacterial diversity in swab specimens (Spearman’s r=0.42, p=0.008; Fig. 3A), however no such correlation was observed in biopsy tissues. No correlation was found between bacterial community and CW duration or ankle-brachial index.

**Figure 3:**
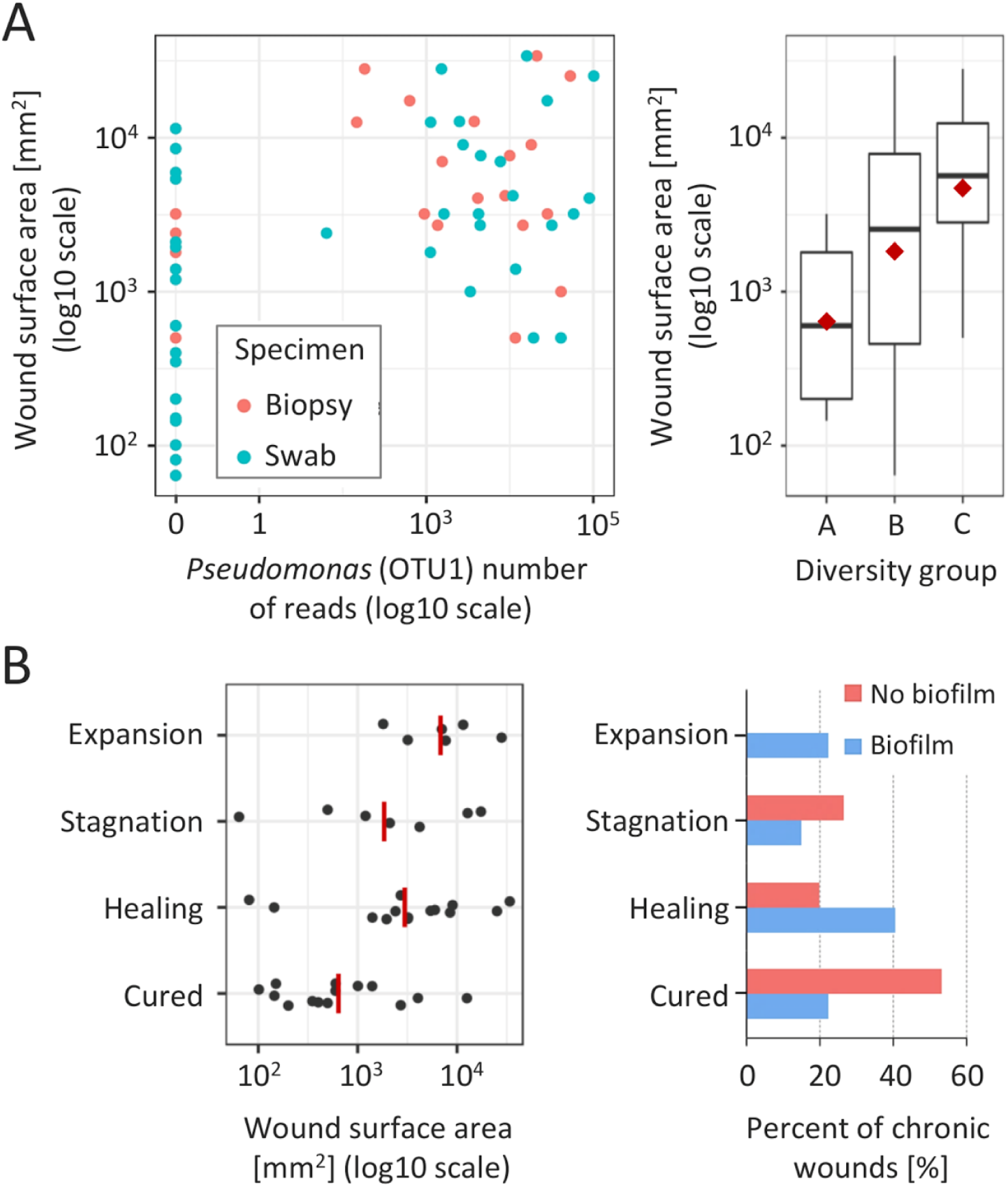
Microbiota association with wound surface area and clinical outcome at one-year follow-up. **(A)** CW surface area was positively correlated with the *Pseudomonas* (OTU1) abundance (left), significant for both biopsy (blue) and swab specimens (red); and with a larger diversity of the bacterial community (right). **(B)** Unfavourable clinical outcomes were most significantly associated with a larger CW surface area (left) and the clinically estimated presence of biofilm (right). Red lines/points denote group means.

CWs were evaluated approximately one year after the sampling and were classified as cured, healing, stagnating or expanding. Higher abundance of *Corynebacterium* (OTU37 and OTU11) was associated with unfavourable clinical outcomes at the follow-ups, most commonly resulting in the stagnation of the healing process (Fisher exact test, p=0.092 and 0.072 for biopsy and swab specimen, respectively). Unfavourable clinical outcomes were also correlated with a larger CW surface area (Spearman’s r=0.44, p=0.004) and the presence of biofilm at the time of CW sampling (Fig. 3B). In the group of CWs with no clinical signs of biofilm, the cure rate was 53.3% (8/15) compared to 22.2% (6/27) cure rate of CWs with clinical signs of biofilm (Fisher exact test, p=0.085). Additionally, expansion of CW area on the follow-up was observed exclusively in cases with estimated biofilm at the first visit (6/45 CWs; 13.3%). In CWs with clinically estimated biofilm, microbiota showed slightly lower richness (3.2 OTUs per sample), compared to biofilm-free CWs (5.3 OTUs per sample) (p=0.027); however, no specific bacterial groups could be correlated to the biofilm formation.

Additionally, we observed a strong positive correlation between five OTUs comprising *Peptoniphilus* (OTU14 and OTU33), *Finegoldia* (OTU18) and *Anaerococcus* (OTU21 and OTU24) (mean Spearman’s r=0.59). At least two of these five OTUs co-occurred in 15/45 CWs (33.3%) and all five co-occurred in 3/45 samples (6.7%) (Fig. S4).

## Discussion

The aim of the present study was to compare different approaches for characterization of bacterial communities residing in CWs. We evaluated the differences between swabs versus biopsy specimens and differences between performing 16S amplicon sequencing (16S-AS) versus cultivation-based methods. Finally, we associated microbiota signatures with patient- and wound-specific parameters and clinical outcomes at the one-year follow-up.

Comparison between swabs and tissue samples did not reveal significant advantage of one method of sampling over the other, which is in concordance with previously reported studies (20, 21). However, we observed that the rate of concordance between cultivation and 16S-AS was higher when analysing biopsy tissue (36.4%) compared to the swab samples (28.7%). Concordance between the swab/biopsy pairs of the same CW was 58.4% and 25.0% for cultivation-based and 16S-AS analysis, respectively. The lower matching rate in case of 16S-AS approach was most likely a consequence of the detection of non-viable bacteria and bacteria present at low cell concentration, which are more prone to the sampling bias.

Previous studies already demonstrated the advantages of using sequencing-based methods over cultivation for the characterization of bacterial communities in CWs (17, 19). In this study, the 16S-AS approach appeared advantageous over cultivation when characterizing communities with a high bacterial diversity. In up to 60% of CWs with the highest bacterial diversity (more than 4 OTUs presented 99% of total obtained sequencing reads), the respective genera was detected solely with 16S-AS. On the other hand, in the samples dominated by a single bacterial group we did not observe any advantage of one method over the other. The majority of bacterial groups that we failed to detect with cultivation, included obligate and facultative anaerobic bacteria from genera *Anaerococcus, Finegoldia, Porphyromonas, Morganella* and *Providencia*, which is in concordance with previous publications (17, 19). Interestingly, the five OTUs corresponding to three Gram-positive anaerobic cocci (*Anaerococcus, Finegoldia* and *Peptoniphilus*) frequently co-occurred in our study, forming a consortium which has already been reported (30), however its clinical relevance remains to be elucidated.

Biofilms, which are present in at least 78% of CWs (31), have prominent influence on development of CWs and hamper CW healing due to the increased resistance against the host defences and antimicrobial therapy (10, 11). In our study, CWs with clinically assessed biofilm showed 31% lower cure rate compared to those without estimated presence of biofilm. CWs with clinically assessed biofilms were associated with a larger surface area, which is well known predictor of impaired CW healing (32, 33). Additionally, all the CWs, which showed the surface area expansion at one-year follow-up, had a diagnosed biofilm at the first visit. In this study, only 15% of CWs with a surface area larger than 1000 mm^2^ healed in a period of one year. Microbiota association analysis revealed that larger CW surface area positively correlated with higher bacterial diversity and increased abundance of *P*. *aeruginosa*, which is in a partial disagreement with previous study by Loesche et al. (7), which reported improved healing of diabetic foot ulcers with higher bacterial diversity.

Increased abundance of *Corynebacterium* was the single most predictive bacterial marker associated with unfavourable clinical outcomes in our study. *Corynebacterium* species are generally perceived as commensal (34); and the abundance of *Corynebacterium* was shown to be inversely correlated with *S*. *aureus*, suggesting potential protective role (35). However, under the right circumstances, *Corynebacterium* species can be clinically relevant (36–38) and the targeted treatment against this bacterium has shown improvement in the CW healing process (39). In this study, only *Corynebacteirum striatum* was identified by cultivation, corresponding to the more prevalent OTU11 in the 16S-AS dataset. However, an additional less prevalent *Corynebacteirum* OTU37 was also associated with unfavourable clinical outcomes. Top BLAST hits suggested either *C*. *pseudodiphtheriticum* or *C*. *propinquum*. Reports on the clinical relevance of these two *Corynebacterium* species in the context of CW infections are rare (40, 41). *Staphylococcus* and *Pseudomonas* have been previously reported as the most prevalent genera present in the microbiota of CWs (6, 42), however no correlation of these two species with CW-specific factors or clinical outcome were found in our dataset.

The comparative analysis of 45 CWs included in this study showed that swabs and biopsy tissues are comparably sufficient for the correct identification of the dominating bacterial colonizers in CWs. Sequencing based approach was more efficient at capturing the broad spectrum of bacteria in communities with high bacterial diversity, detecting multiple additional obligate and facultative anaerobic bacterial taxa. Two *Corynebacterium* species were found to be associated with unfavourable clinical outcome.

## Acknowledgements

This work was in part supported by Slovenian Research Agency (grant J2-7413). Authors would like to acknowledge the contributions by Slavica Lorencic Robnik, Matija Primec and Aleksander Kocuvan.

## References

1. Grey JE, Enoch S, Harding KG. 2006. Wound assessment. BMJ 332:285–288.

2. Mekkes JR, Loots M a. M, Van Der Wal AC, Bos JD. 2003. Causes, investigation and treatment of leg ulceration. Br J Dermatol 148:388–401.

3. Martinengo L, Olsson M, Bajpai R, Soljak M, Upton Z, Schmidtchen A, Car J, Järbrink K. 2019. Prevalence of chronic wounds in the general population: systematic review and meta-analysis of observational studies. Ann Epidemiol 29:8–15.

4. Schultz G, Bjarnsholt T, James GA, Leaper DJ, McBain AJ, Malone M, Stoodley P, Swanson T, Tachi M, Wolcott RD. 2017. Consensus guidelines for the identification and treatment of biofilms in chronic nonhealing wounds. Wound Repair Regen 25:744–757.

5. Kalan L, Loesche M, Hodkinson BP, Heilmann K, Ruthel G, Gardner SE, Grice EA. 2016. Redefining the Chronic-Wound Microbiome: Fungal Communities Are Prevalent, Dynamic, and Associated with Delayed Healing. mBio 7: e01058–16.

6. Wolcott RD, Hanson JD, Rees EJ, Koenig LD, Phillips CD, Wolcott RA, Cox SB, White JS. 2016. Analysis of the chronic wound microbiota of 2,963 patients by 16S rDNA pyrosequencing. Wound Repair Regen 24:163–174.

7. Loesche M, Gardner SE, Kalan L, Horwinski J, Zheng Q, Hodkinson BP, Tyldsley AS, Franciscus CL, Hillis SL, Mehta S, Margolis DJ, Grice EA. 2017. Temporal Stability in Chronic Wound Microbiota Is Associated With Poor Healing. J Invest Dermatol 137:237–244.

8. Bartow-McKenney C, Hannigan GD, Horwinski J, Hesketh P, Horan AD, Mehta S, Grice EA. 2018. The microbiota of traumatic, open fracture wounds is associated with mechanism of injury. Wound Repair Regen 26:127–135.

9. Kalan LR, Meisel JS, Loesche MA, Horwinski J, Soaita I, Chen X, Uberoi A, Gardner SE, Grice EA. 2019. Strain-and Species-Level Variation in the Microbiome of Diabetic Wounds Is Associated with Clinical Outcomes and Therapeutic Efficacy. Cell Host Microbe 25:641–655.

10. Percival SL, McCarty SM, Lipsky B. 2015. Biofilms and Wounds: An Overview of the Evidence. Adv Wound Care 4:373–381.

11. Metcalf DG, Bowler PG. 2015. Biofilm delays wound healing: A review of the evidence. Burns Trauma 1:5–12.

12. Tuttle MS. 2015. Association Between Microbial Bioburden and Healing Outcomes in Venous Leg Ulcers: A Review of the Evidence. Adv Wound Care 4:1–11.

13. O’Meara S, Al-Kurdi D, Ologun Y, Ovington LG. 2010. Antibiotics and antiseptics for venous leg ulcers. Cochrane Database Syst Rev CD003557.

14. Haalboom M, Blokhuis-Arkes MHE, Beuk RJ, Meerwaldt R, Klont R, Schijffelen MJ, Bowler PB, Burnet M, Sigl E, van der Palen J a. M. 2019. Culture results from wound biopsy versus wound swab: does it matter for the assessment of wound infection? Clin Microbiol Infect 25:629.e7-629.e12.

15. Gu W, Miller S, Chiu CY. 2019. Clinical Metagenomic Next-Generation Sequencing for Pathogen Detection. Annu Rev Pathol 14:319–338.

16. Tatum OL, Dowd SE. 2012. Wound Healing Finally Enters the Age of Molecular Diagnostic Medicine. Adv Wound Care 1:115–119.

17. Smith K, Collier A, Townsend EM, O’Donnell LE, Bal AM, Butcher J, Mackay WG, Ramage G, Williams C. 2016. One step closer to understanding the role of bacteria in diabetic foot ulcers: characterising the microbiome of ulcers. BMC Microbiol 16:54.

18. Malone M, Johani K, Jensen SO, Gosbell IB, Dickson HG, Hu H, Vickery K. 2017. Next Generation DNA Sequencing of Tissues from Infected Diabetic Foot Ulcers. EBioMedicine 21:142–149.

19. Boers SA, Hiltemann SD, Stubbs AP, Jansen R, Hays JP. 2018. Development and evaluation of a culture-free microbiota profiling platform (MYcrobiota) for clinical diagnostics. Eur J Clin Microbiol Infect Dis 37:1081–1089.

20. Gardiner M, Vicaretti M, Sparks J, Bansal S, Bush S, Liu M, Darling A, Harry E, Burke CM. 2017. A longitudinal study of the diabetic skin and wound microbiome. PeerJ 5:e3543.

21. Slater RA, Lazarovitch T, Boldur I, Ramot Y, Buchs A, Weiss M, Hindi A, Rapoport MJ. 2004. Swab cultures accurately identify bacterial pathogens in diabetic foot wounds not involving bone. Diabet Med 21:705–709.

22. Gardner SE, Frantz RA, Saltzman CL, Hillis SL, Park H, Scherubel M. 2006. Diagnostic validity of three swab techniques for identifying chronic wound infection. Wound Repair Regen 14:548–557.

23. Snyder RJ, Bohn G, Hanft J, Harkless L, Kim P, Lavery L, Schultz G, Wolcott R. 2017. Wound Biofilm: Current Perspectives and Strategies on Biofilm Disruption and Treatments. Wounds 29:S1–S17.

24. Hurlow J, Blanz E, Gaddy JA. 2016. Clinical investigation of biofilm in non-healing wounds by high resolution microscopy techniques. J Wound Care 9:S11–22.

25. Angel DE, Lloyd P, Carville K, Santamaria N. 2011. The clinical efficacy of two semi-quantitative wound-swabbing techniques in identifying the causative organism(s) in infected cutaneous wounds. Int Wound J 8:176–185.

26. Klindworth A, Pruesse E, Schweer T, Peplies J, Quast C, Horn M, Glockner FO. 2013. Evaluation of general 16S ribosomal RNA gene PCR primers for classical and next-generation sequencing-based diversity studies. Nucleic Acids Res 41:e1.

27. Schloss PD, Westcott SL, Ryabin T, Hall JR, Hartmann M, Hollister EB, Lesniewski RA, Oakley BB, Parks DH, Robinson CJ, Sahl JW, Stres B, Thallinger GG, Horn DJV, Weber CF. 2009. Introducing mothur: Open-Source, Platform-Independent, Community-Supported Software for Describing and Comparing Microbial Communities. Appl Environ Microbiol 75:7537–7541.

28. Kozich JJ, Westcott SL, Baxter NT, Highlander SK, Schloss PD. 2013. Development of a Dual-Index Sequencing Strategy and Curation Pipeline for Analyzing Amplicon Sequence Data on the MiSeq Illumina Sequencing Platform. Appl Environ Microbiol 79:5112–5120.

29. Eren AM, Borisy GG, Huse SM, Welch JLM. 2014. Oligotyping analysis of the human oral microbiome. PNAS 111:E2875–E2884.

30. Choi Y, Banerjee A, McNish S, Couch KS, Torralba MG, Lucas S, Tovchigrechko A, Madupu R, Yooseph S, Nelson KE, Shanmugam VK, Chan AP. 2019. Co-occurrence of Anaerobes in Human Chronic Wounds. Microb Ecol 77:808–820.

31. Malone M, Bjarnsholt T, McBain AJ, James GA, Stoodley P, Leaper D, Tachi M, Schultz G, Swanson T, Wolcott RD. 2017. The prevalence of biofilms in chronic wounds: a systematic review and meta-analysis of published data. J Wound Care 26:20–25.

32. Margolis DJ, Allen-Taylor L, Hoffstad O, Berlin JA. 2002. Diabetic Neuropathic Foot Ulcers: The association of wound size, wound duration, and wound grade on healing. Diabetes Care 25:1835–1839.

33. Kramer JD. 1997. Patient, wound and treatment characteristics associated with healing in pressure ulcers. Adv Skin Wound Care 13:17–24.

34. Scharschmidt TC, Fischbach MA. 2013. What Lives On Our Skin: Ecology, Genomics and Therapeutic Opportunities Of the Skin Microbiome. Drug Discov Today Dis Mech 10:e83–e89.

35. Uehara Y, Nakama H, Agematsu K, Uchida M, Kawakami Y, Abdul Fattah AS, Maruchi N. 2000. Bacterial interference among nasal inhabitants: eradication of Staphylococcus aureus from nasal cavities by artificial implantation of Corynebacterium sp. J Hosp Infect 44:127–133.

36. Leal SM, Jones M, Gilligan PH. 2016. Clinical Significance of Commensal Gram-Positive Rods Routinely Isolated from Patient Samples. J Clin Microbiol 54:2928–2936.

37. Dowd SE, Wolcott RD, Sun Y, McKeehan T, Smith E, Rhoads D. 2008. Polymicrobial Nature of Chronic Diabetic Foot Ulcer Biofilm Infections Determined Using Bacterial Tag Encoded FLX Amplicon Pyrosequencing (bTEFAP). PLoS One 3:e3326.

38. Bessman AN, Geiger PJ, Canawati H. 1992. Prevalence of Corynebacteria in diabetic foot infections. Diabetes Care 15:1531–1533.

39. Wolcott RD, Cox SB, Dowd SE. 2010. Healing and healing rates of chronic wounds in the age of molecular pathogen diagnostics. J Wound Care 19:272–278.

40. Saïdani M, Kammoun S, Boutiba-Ben Boubaker I, Ben Redjeb S. 2010. Corynebacterium propinquum isolated from a pus collection in a patient with an osteosynthesis of the elbow. Tunis Med 88:360–362.

41. Cantarelli VV, Brodt TCZ, Secchi C, Inamine E, Pereira F de S. 2008. Cutaneous infection caused by Corynebacterium pseudodiphtheriticum: a microbiological report. Rev Inst Med Trop Sao Paulo 50:51–52.

42. Dowd SE, Sun Y, Secor PR, Rhoads DD, Wolcott BM, James GA, Wolcott RD. 2008. Survey of bacterial diversity in chronic wounds using Pyrosequencing, DGGE, and full ribosome shotgun sequencing. BMC Microbiology 8:43.

